# Improved Algorithms for Finding Edit Distance Based Motifs

**DOI:** 10.1101/020131

**Authors:** Soumitra Pal, Sanguthevar Rajasekaran

## Abstract

Motif search is an important step in extracting meaningful patterns from biological data. Since the general problem of motif search is intractable, there is a pressing need to develop efficient exact and approximation algorithms to solve this problem. We design novel algorithms for solving the *Edit-distance-based Motif Search (EMS)* problem: given two integers *l, d* and *n* biological strings, find all strings of length *l* that appear in each input strings with at most *d* substitutions, insertions and deletions. These algorithms have been evaluated on several challenging instances. Our algorithm solves a moderately hard instance (11, 3) in a couple of minutes and the next difficult instance (14, 3) in a couple of hours whereas the best previously known algorithm, EMS1, solves (11, 3) in a few hours and does not solve (13, 4) even after 3 days. This significant improvement is due to a novel and provably efficient neighborhood generation technique introduced in this paper. This efficient approach can be used in other edit distance based applications in Bioinformatics, such as *k*-spectrum based sequence error correction algorithms. We also use a trie based data structure to efficiently store the candidate motifs in the neighbourhood and to output the motifs in a sorted order.

## Introduction

Motif search has applications in solving such crucial problems as identification of alternative splicing sites, determination of open reading frames, identification of promoter elements of genes, identification of transcription factors and their binding sites etc. (see e.g., Nicolae and Rajasekaran^1^). There are many formulations of the motif search problem. A widely studied formulation is known as (*l, d*)-motif search or Planted Motif Search (PMS).^2^ Given two integers *l, d* and *n* biological strings the problem is to find all short strings of length *l* that appear in each of the *n* input strings with at most *d* mismatches. There is a significant amount of work in the literature on PMS [1, 3, 4, and so on].

PMS considers only point mutations as events of divergence in biological sequences. However, insertions and deletions also play important roles in divergence.^2, 5^ Therefore, researchers have also considered a formulation in which the Levenshtein distance (or edit distance), instead of mismatches, is used for measuring the degree of divergence.^6, 7^ Given *n* strings *S*^(1)^, *S*^(2)^, *…, S*^(*n*)^, each of length *m* from a fixed alphabet Σ, and integers *l, d*, the Edit-distance-based Motif Search (EMS) problem is to find all patterns *M* of length *l* that occur in at least one position in each *S*^(*i*)^ with an edit distance of at most *d*. More formally, *M* is a motif if and only if *∀i*, there exist *k* ∈ [*l - d, l* + *d*], *j* ∈ [1, *m - k* + 1] such that for the substring 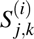 of length *k* at position *j* of *S*^(*i*)^, 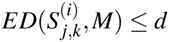. Here *ED*(*X, Y*) stands for the edit distance between two strings *X* and *Y*.

EMS is also NP-hard since PMS is a special case of EMS and PMS is known to be NP-hard.^8^ As a result, any exact algorithm for EMS that finds all the motifs for a given input can be expected to have an exponential (in some of the parameters) worst case runtime. One of the earliest EMS algorithms is due to Rocke and Tompa^6^ and is based on Gibbs Sampling which requires repeated searching of the motifs in a constantly evolving collection of aligned strings, and each search pass requires *O*(*nl*) time. Sagot^7^ gave a suffix tree based algorithm that takes *O*(*n*^2^*ml^d^ |*Σ*|* ^*d*^) time and *O*(*n*^2^*m/w*) space where *w* is the word length of the computer. Adebiyi and Kaufmann^9^ proposed an algorithm with an expected runtime of *O*(*nm* + *d*(*nm*)^(1+*pow*(*ε*))^ log *nm*) where *ε* = *d/l* and *pow*(*ε*) is an increasing concave function. The value of *pow*(*ε*) is roughly 0.9 for protein and DNA sequences.

Rajasekaran et al.^10^ proposed a simpler Deterministic Motif Search (DMS) that has the same worst case time complexity as the algorithm by Sagot.^7^ The algorithm generates and stores the neighbourhood of every substring of length in the range [*l - d, l* + *d*] of every input string and using a radix sort based method outputs the neighbours that are common to at least one substring of each input string. This algorithm was implemented by Pathak et al.^11^

Following a useful practice for PMS algorithms, Pathak et al.^11^ evaluated their algorithm on certain instances that are considered challenging and are generated as follows: *n* = 20 random DNA/protein strings of length *m* = 600, and a short random string *M* of length *l* are generated according to the independent identically distributed (i.i.d) model. A separate random *d*-edit distance neighbour of *M* is “planted” in each of the *n* input strings. Such an (*l, d*) instance is defined to be a challenging instance if *d* is the largest integer for which the expected number of spurious motifs, i.e., the motifs of length *l* that would occur in the input by random chance, does not exceed a constant (10000 in this paper).

Table 4 shows the expected number of spurious motifs in the random instances for *l* in the range [5, 17] and for *d* in the range [0, *l* - 2] computed using equation (27) given in Appendix. Though the expected number of spurious motifs in EMS instances are in general different from those in PMS instances, it turns out that the challenging instances are the same for both problems for the range of *l, d* we considered: (7, 1), (9, 2), (11, 3), (13, 4), and (15, 5).

The algorithm by Pathak et al.^11^ solves the moderately hard instance (11, 3) in a few hours and does not solve the next difficult instance (13, 4) even after 3 days. A key time consuming part of the algorithm is in the generation of the edit distance neighbourhood of all subsequences as there are many common neighbours.

## Contributions

In this paper we present an improved algorithm for EMS that solves the instance (11, 3) in less than a couple of minutes and the instance (14, 3) in less than a couple of hours. Our algorithm uses an efficient technique (introduced in this paper) to generate the edit distance neighbourhood of length *l* with distance at most *d* of all substrings of an input string.

We prove that every substring identified by our neighborhood generation technique as a neighbor of a string is nearly distinct. In other words, our neighborhood generation technique does not spend a lot of time generating neighbors that have already been generated. This efficient approach can be used in other edit distance based applications in Bioinformatics, such as generating the candidate set of corrections in *k*-spectrum based error correction algorithms^12^ for sequences with insertion and deletion errors.

We use a trie based data structure to efficiently store the neighbourhood. This not only simplifies the removal of duplicate neighbours but also helps in outputting the final motifs in sorted order using a depth first search traversal of the trie.

A C++ implementation of our algorithm is available at 

~~~
https://github.com/soumitrakp/ems2.git
~~~

.

## Methods

In this section we introduce some notations and observations.

An (*l, d*)*-friend* of a *k*-mer *L* is an *l*-mer that is at an exact distance of *d* from *L*. Let *F*_*l,d*_ (*L*) denote the set of all (*l, d*)-friends of *L*. An (*l, d*)*-neighbour* of a *k*-mer *L* is an *l*-mer that is at a distance of no more than *d* from *L*. Let *N*_*l,d*_ (*L*) denote the set of all (*l, d*)-neighbours of *L*. Then

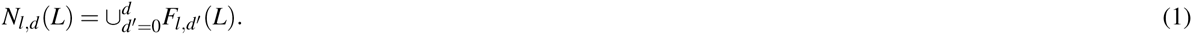

For a string *S* of length *m*, an (*l, d*)*-motif* of *S* is an *l*-mer that is at a distance no more than *d* from some substring of *S*. Thus an (*l, d*)-motif of *S* is an (*l, d*)-neighbour of at least one substring *S*_*j,k*_ = *S*_*j*_*S*_*j*+1_ … *S*_*j+k*-1_ where *k* ∈ [*l - d, l* + *d*]. Therefore, the set of (*l, d*)-motifs of *S*, denoted by *M*_*l,d*_ (*S*), is given by

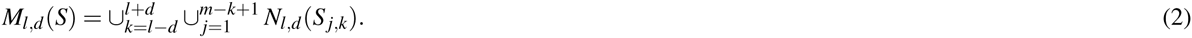

For a collection of strings *𝒮* = {*S*^(1)^, *S*^(2)^, …, *S*^(*m*)^}, a (common) (*l, d*)-motif is an *l*-mer that is at a distance no more than *d* from at least one substring of each *S*^(*i*)^. Thus the set of (common) (*l, d*)-motifs of *𝒮*, denoted by *M*_*l,d*_ (*𝒮*), is given by

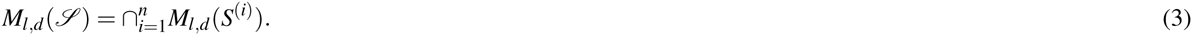

One simple way of computing *F*_*l,d*_ (*L*) is to grow the friendhood of *L* by one distance at a time for *d* times and to select only the friends having length *l*. Let *G*(*L*) denote the set of strings obtained by one edit operation on *L* and 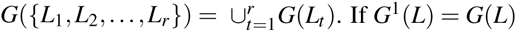, and for *t* > 1, *G*^*t*^(*L*) = *G*(*G*^*t*-1^(*L*)) then

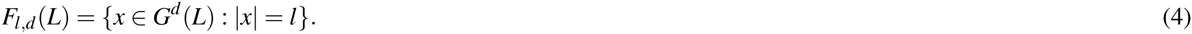

Using equations (1), (2), (3) and (4), Pathak et al.^11^ gave an algorithm that stores all possible candidate motifs in an array of size |Σ|^*l*^. However the algorithm is inefficient in generating the neighbourhood as the same candidate motif is generated by several combinations of the basic edit operations. Also, the *O*(|Σ|^*l*^) memory requirement makes the algorithm inapplicable for larger instances. In this paper we mitigate these two limitations.

## Efficient Neighbourhood Generation

We now give a more efficient algorithm to generate the (*l, d*)-neighbourhood of all possible *k*-mers of a string. Instead of computing (*l, d′*)-friendhood for all 0 ≤ *d*′ ≤ *d*, we compute only the exact (*l, d*)-friendhood.

### Lemma 1.

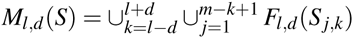.

*Proof.* Consider the *k*-mer *L* = *S*_*j,k*_. If *k* = *l - d* then we need *d* insertions to make *L* an *l*-mer. There cannot be any (*l, d′*)-neighbour of *L* for *d*′ < *d*. Thus

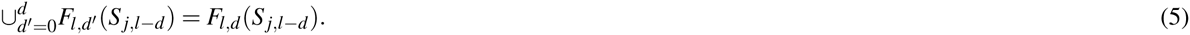

Suppose *k* > *l - d*. Any (*l, d -* 1)-neighbour *B* of *L* is also an (*l, d*)-neighbour of *L*′ = *S*_*j,k-*1_ because *ED*(*B, L*′) ≤ *ED*(*B, L*) + *ED*(*L, L′*) ≤ (*d -* 1) + 1 = *d*. Thus

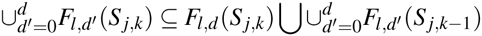

which implies that

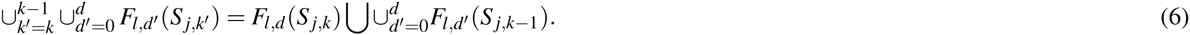

Applying equation (6) repeatedly for *k* = *l* + *d, l* + *d* 1, *…, l − d* + 1, along with equation (5) in equations (1) and (2) gives the result of the lemma. □

We generate *F*_*l,d*_ (*S*_*j,k*_) in three phases: we apply *δ* deletions in the first phase, *β* substitutions in the second phase, and finally *α* insertions in the final phase, where *d* = *δ* + *α* + *β* and *l* = *k - δ* + *α*. Solving for *α, β, δ* gives max {0, *q*} ≤ *δ ≤* (*d* + *q*)*/*2, *α* = *δ - q* and *β* = *d - α - δ* = *d -* 2*δ* + *q* where *q* = *k - l, q* ∈ [*-d,* +*d*]. In each of the phases, the neighbourhood is grown by one edit operation at a time. Duplication in the friendhood is avoided as follows: if the current edit operation is at position *j*, the subsequent edit operation in the same phase are restricted at positions larger than *j*. Formally,

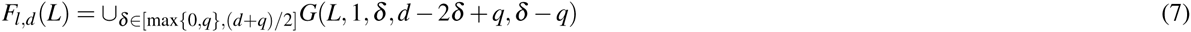

where *q* = |*L*| − *l* and *G*(*L, j, δ, β, α*)

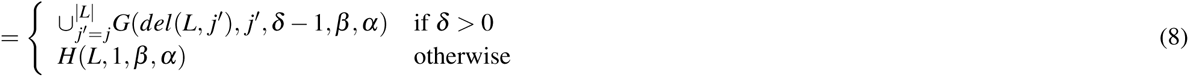

where *del*(*L, j*) is the string obtained by deleting *L*_*j*_ and *H*(*L, j, β, α*) =

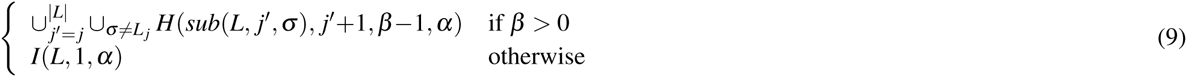

where *sub*(*L, j, σ*) is obtained by substituting *L*_*j*_ by *σ* and *I*(*L, j, α*)

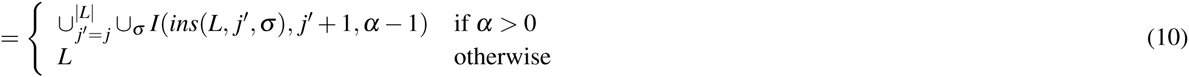

where *ins*(*L, j, σ*) is obtained by inserting *σ* just before *L*_*j*_.

## Compact Motifs

Let us consider an example 3-mer *σ*_1_*σ*_2_*σ*_3_ and all possible single insertions just before position 2. The |Σ| neighbours thus generated can be compactly represented as *σ*_1_*\*σ*_2_*σ*_3_ where represents any character in Σ. Similarly a substitution at position 2 can be compactly represented by *σ*_1_*\*σ*_3_ where *\** represents any character in Σ \ *σ*_2_. This can be generalized for any number of substitutions and insertions.

Let Σ*′* represent the compact alphabet Σ ∪ {*} where * is a special character not included in Σ. The *expansion* of a compact string *L* ∈ Σ*′*^+^ is defined as *X*(*L*) = {*T* ∈ Σ^|*L*|^ | *T*_*j*_ ≠ *L*_*j*_ ⇒ *L*_*j*_ = *}. The representation of a set of strings in Σ^+^ using compact strings in Σ*′*^+^ is not unique. For example, if Σ = {0, 1}, the set {000, 001, 100, 101} can be represented by {00*\*,* 100, 101}, {00*\*,* 10*}, {*0*}, and so on. Here we are interested in only those compact strings that arise if we use the special character for insertion and substitution. Let the compact friendhood thus generated be denoted as 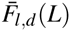 and formally defined as

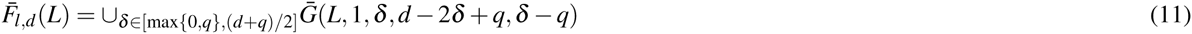

where *q* = |*L*| - *l* and 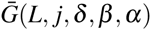

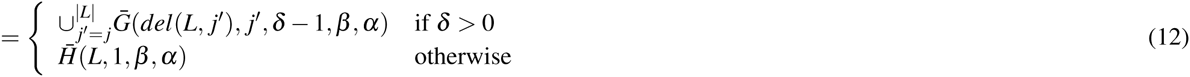

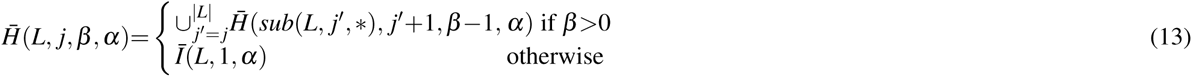

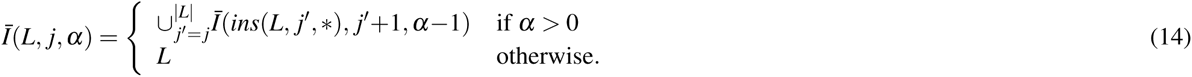

For *𝒮* = {*S*^(1)^, *S*^(2)^, …, *S*^(*m*)^}, let

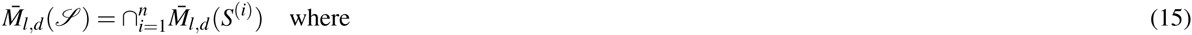

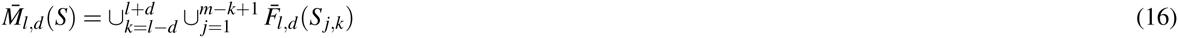

with the usual definition of union and the following definition of intersection on compact strings *A, B* ∈ Σ′^*l*^

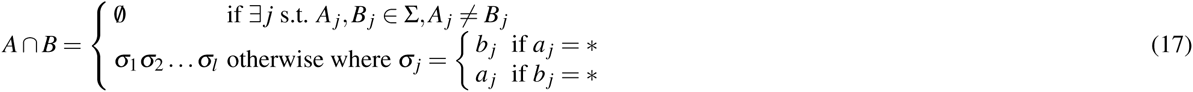

and on sets of compact strings in Σ′^*l*^

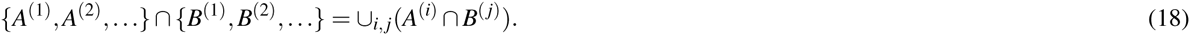

### Lemma 2.

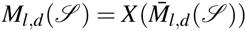.

*Proof.* The equivalence is straight forward if there are only insertions. For substitution too the equivalence is valid as *L* is already in the neighbourhood and 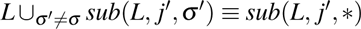. □

## Trie for Storing Compact Motifs

We store the compact motifs 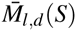 in a trie based data structure which we call a *motif trie*. This helps implement the intersection defined in equation (17) and required in equation (15). Each node in the motif trie has at most |Σ| children. The edges from a node *u* to its children *v* are labeled with mutually exclusive subsets *label*(*u, v*) *⊆* Σ. An empty set of compact motifs is represented by a single root node. A non-empty trie has *l* + 1 levels of nodes, the root being at level 0. The trie stores the *l*-mer *σ*_1_*σ*_2_ … *σ*_*l*_, all *σ*_*j*_ ∈ Σ, if there is a path from the root to a leaf where *σ _j_* appears in the label of the edge from level *j -* 1 to level *j*.

For each string *S* = *𝒮*^(*i*)^ we keep a separate motif trie *M*^(*i*)^. Each compact neighbour 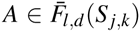 generated by equation (11) is inserted into the motif trie recursively as follows. We start with the root node where we insert *A*_1_*A*_2_ … *A*_*l*_. At a node *u* at level *j* where the prefix *A*_1_*A*_2_ … *A*_*j*-1_ is already inserted, we insert the suffix *A*_*j*_*A*_*j*+1_ … *A*_*l*_ as follows. If *A*_*j*_ ∈ Σ we insert *A*′ = *A* _*j*+1_*A* _*j*+2_ … *A*_*l*_ to the children *v* of *u* such that *A*_*j*_ ∈ *label*(*u, v*). If *label*(*u, v*) {*A*_*j*_}, before inserting we make a copy of sub trie rooted at *v*. Let *v*′ be the root of the new copy. We make *v*′ a new child of *u*, set *label*(*u, v′*) = {*A*_*j*_}, remove *A*_*j*_ from *label*(*u, v*), and insert *A*′ to *v′*. On the other hand if *A*_*j*_ = we insert *A*′ to each children of *u*. Let *T* = Σ if *A*_*j*_ = and *T* = {*A*_*j*_} otherwise. Let *R* = *T ∪_v_label*(*u, v*). If *T* ≠ Ø we create a new child *v*′ of *u*, set *label*(*u, v′*) = *R* and recursively insert *A*′ to *v′*. Fig. 1 shows examples of inserting into the motif trie.

**Figure 1.**
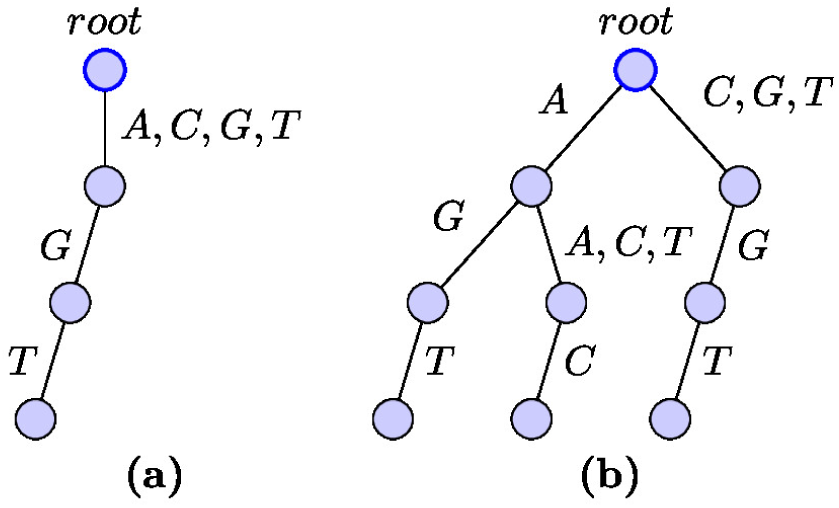
Inserting into motif trie for Σ = {*A, C, G, T*} and *l* = 2. (a) After inserting *\*GT* into empty trie. (b) After inserting another string *A*C*.

We also maintain a motif trie for the *ℳ* for the common compact motifs found so far, starting with *ℳ* = *M*^(1)^. After processing string *S*^(*i*)^ we intersect the root of *M*^(*i*)^ with the root of *ℳ*. In general a node *u*_2_ ∈ *M*^(*i*)^ at level *j* is intersected with a node *u*_1_ ∈ *ℳ* at level *j* using the procedure shown in Algorithm 1. Fig. 2 shows an example of the intersection of two motif tries.

**Figure 2.**
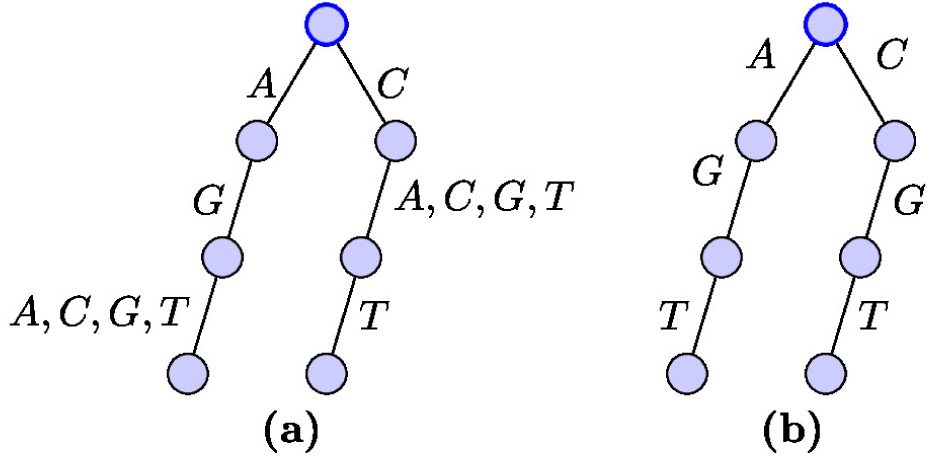
Intersection of motif tries. (a) Trie for *AG* ∪C*T*. (b) Intersection of trie in Fig. 1(b) and trie in Fig. 2(a).

The final set of motifs 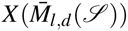 is obtained by a depth-first traversal of *ℳ* outputting the label of the path from the root whenever a leaf is traversed. An edge (*u, v*) is traversed separately for each *σ* ∈ *label*(*u, v*).

**Algorithm 1:**
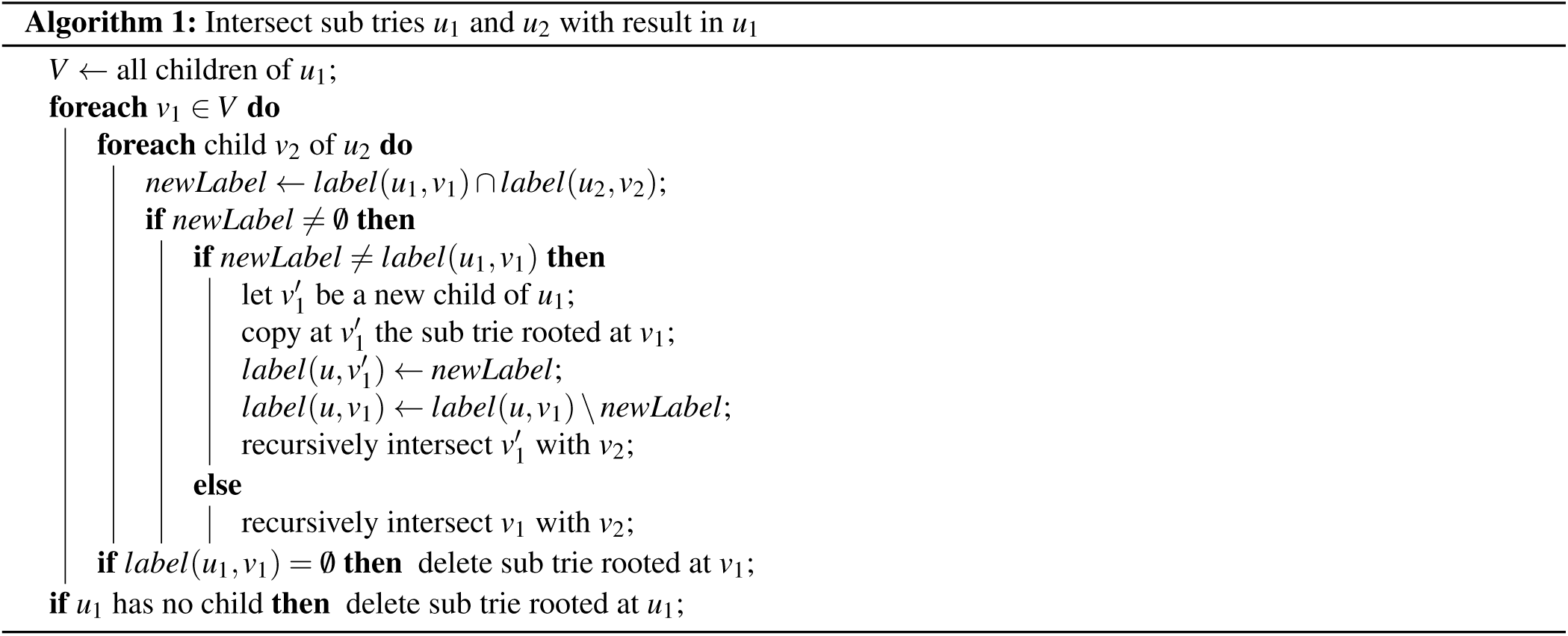
Intersect sub tries *u*_1_ and *u*_2_ with result in *u*_1_

## Efficient Compact Neighbourhood Generation

The set of compact motifs 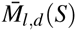 computed in equation (15) is in fact a multiset as the compact motifs generated are not all distinct. A significant part of the time taken by our algorithm is in inserting compact neighbours into the motif trie as it is executed for each neighbour in the friendhood. We improve the performance of our algorithm using some simple rules to reduce the number of times a compact motif is generated. Later we will see that these rules are quite close to the ideal as we will prove that the compact motif generated after skipping using the rules, are distinct if all the characters in the string are distinct.

To differentiate multiple instances of the same element in the multiset 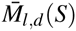 we augment it with the information about how it is generated by equation (15). Formally, each instance *L* of an element in 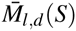 is represented as an ordered tuple 〈*M, S*_*j,k*_, *T*〉 where the sequence of edit operations *T* when applied to *S*_*j,k*_ gives *M*. Each edit operation in *T* is represented as a tuple 〈*p, t*〉 where *p* denotes the position in *S* where the edit operation is applied and *t* ∈ {*D, R, I*} denotes the type of the operation - deletion, substitution and insertion, respectively. At each position there can be one deletion or one substitution but one or more insertions. The tuples in *T* are sorted lexicographically with the normal order for *p* and for *t*, *D* < *R* < *I*. In the rest of the paper we assume that 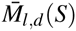 is a set of tuples 〈*M, S*_*j,k*_, *T*〉.

The rules for skipping the elements of the multiset 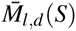 are shown in Table 1. We first show that the rules do not exclude any necessary compact motif. Let 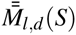 be the set of compact motifs generated by equation (15) except those excluded by Rules 1-9. Let Γ(〈*M, S*_*j,k*_, *T*〉) = *M* and Γ(*Z*) = ∪_*L*∈*Z*_ Γ(*L*).

**Table 1.** Conditions for skipping compact motif *L* = 〈*M, S*_*j,k*_, *T*〉

| Rule | Conditions (in all rules $t \geq 0$) |
| --- | --- |
| 1 | $(j+k \leq m) \wedge \langle j, D \rangle \in T$ |
| 2 | $\{\langle j+t, D \rangle, \langle j+t+1, R \rangle\} \subseteq T$ |
| 3 | $(j+k \leq m) \wedge \{\langle j, R \rangle, \langle j+1, R \rangle, \dots, \langle j+t, R \rangle, \langle j+t+1, D \rangle\} \subseteq T$ |
| 4 | $\{\langle j+t, D \rangle, \langle j+t, I \rangle\} \subseteq T$ |
| 5 | $\{\langle j+t, D \rangle, \langle j+t+1, I \rangle\} \subseteq T$ |
| 6 | $\{\langle j+t, R \rangle, \langle j+t, I \rangle\} \subseteq T$ |
| 7 | $(j > 1) \wedge \langle j, I \rangle \in T$ |
| 8 | $(j > 1) \wedge \{\langle j, R \rangle, \langle j+1, R \rangle, \dots, \langle j+t, R \rangle, \langle j+t+1, I \rangle\} \subseteq T$ |
| 9 | $(j+k \leq m) \wedge \langle j+k, I \rangle \in T$ |

### Lemma 3.

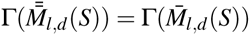.

*Proof.* By construction, 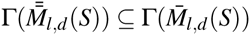. We show by contradiction that 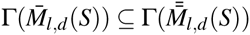.

Let 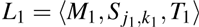 and 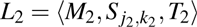 be two elements of 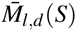 and 〈*p*_1_, *t*_1_〉 ∈ *T*_1_, 〈*p*_2_, *t*_2_〉 ∈ *T*_2_ be the leftmost edit operations where *T*_1_, *T*_2_ differ. We impose an order *L*_1_ < *L*_2_ if and only if (*k*_1_ < *k*_2_) ∨ ((*k*_1_ = *k*_2_) ∧ (*p*_1_ < *p*_2_)) ∨ ((*k*_1_ = *k*_2_) ∧ (*p*_1_ = *p*_2_) ∧ (*t*_1_ < *t*_2_)).

Let *L* = 〈*M, S*_*j,k*_, *T*〉 be the largest (in the order defined above) element in 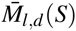 such that there is no element 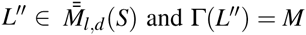. For each Rule 1-9 we show a contradiction that if *L* is skipped by the rule then there is another 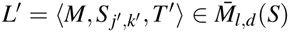 with the same number of edit operations but *L* < *L*′. Fig. 3 illustrates the choice of *L*′ under different rules.

**Figure 3.**
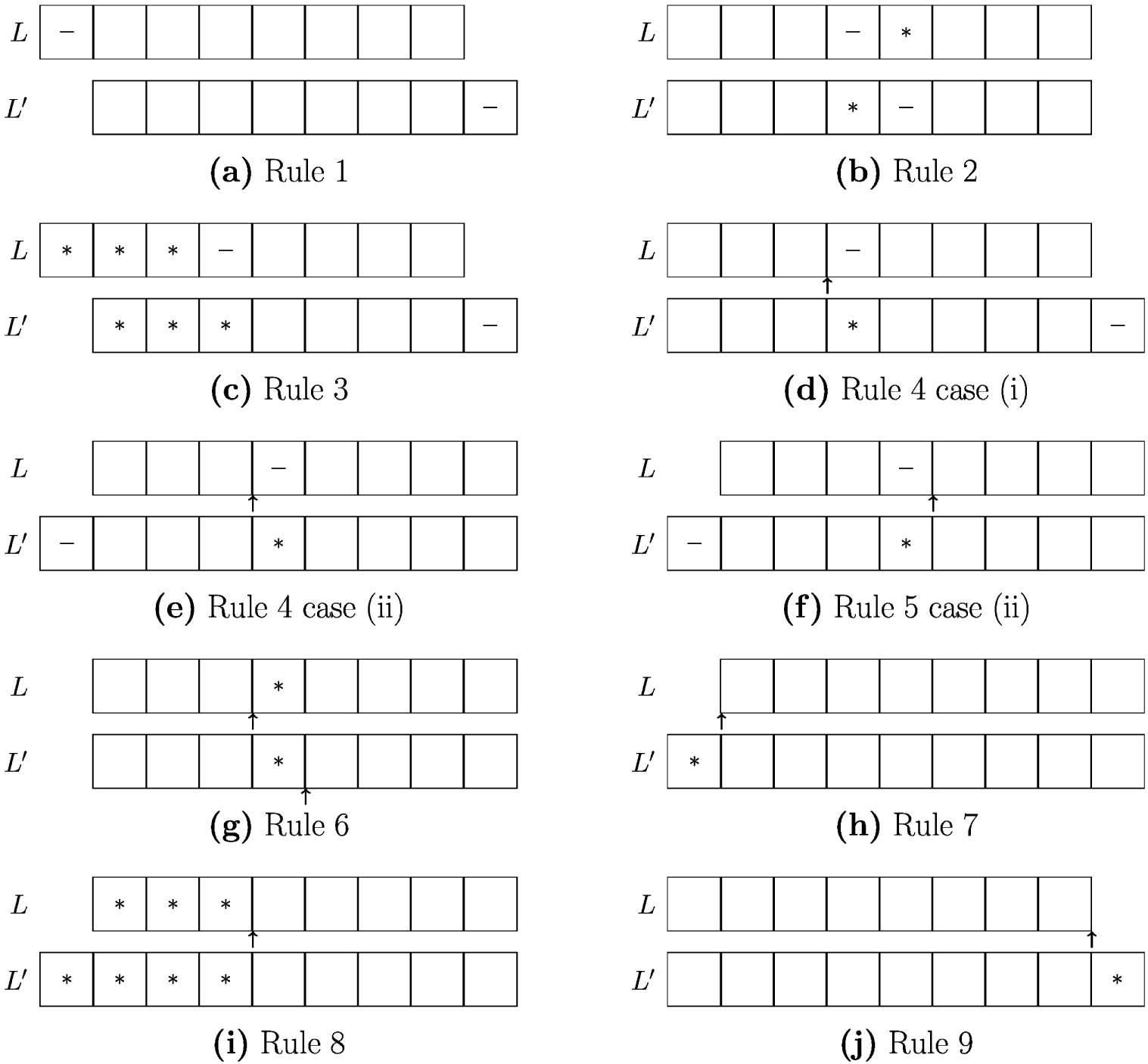
Construction of *L*^*′*^ under different rules in the proof of Lemma 3. Insertions are shown using arrows, deletions using *-* and substitutions using *. Rule 5 case (i) is similar to Rule 4 case (i) and omitted to save space.

Rule 1. Here *j* + *k* ≤ *m* and 〈*j, D*〉 ∈ *T*. Consider *T*′ = *T* \ 〈*j, D*〉) ∪ 〈*j + k*, *D*〉, and *j*′ = *j* + 1, *k*′ = *k*.

Rule 2. Consider *T*′ = *T* \ (〈*j* + *t*, *D*〉, 〈*j + t* + 1, *R*〉} ∪ {〈*j + t*, *R*〉, 〈*j + t* + 1, *D*〉}, and *j*′ = *j*,*k*′ = *k*.

Rule 3. *T*′ = *T*\{〈*j,R*〉, 〈*j+t*+1,*D*〉} ∪ {〈*j+t*+1, *R*〉, 〈*j+k*,*D*〉}, *j*′ = *j* + 1,*k*′ = *k*.

Rule 4. For this and subsequent rules *k* < *l + d* as there is at least one insertion and hence *k*′ could possibly be equal to *k* + 1. We consider two cases. Case (i) *j + k* ≤ *m*: *T*′ = *T*\{〈*j +t*,*D*〉, 〈*j +t,I*〉} ∪ {〈*j+t,R*〉, 〈*j + k,D*〉}, *j*′ = *j,k*′ = *k* + 1. Case (ii) *j* + *k* = *m* + 1: Here deletion of *S*_*j*_ is allowed by Rule 1. *T*′ = *T* \ {(*j + t,D*}, (*j +t,I*}} ∪ {(*j - 1,D*}, (*j+t,R*}}, *j*′ = *j* - 1, *k*′ = *k* + 1.

Rule 5. The same argument for Rule 4 applies considering 〈*j+t*+1,*I*〉 instead of 〈*j+t*, *I*〉.

Rule 6. *T*′ = *T* \ 〈*j+t, I*〉 ∪ 〈*j+t+1,I*〉, and *j*′ = *j, k*′ = *k*.

Rule 7. *T*′ = *T* \ 〈*j, I*〉 ∪ 〈*j*-1,*R*〉, *j*′ = *j* - 1, *k*′ = *k* + 1.

Rule 8. *T*′ = *T* \ 〈*j+t*, *I*〉 ∪ 〈*j*-1,*R*〉, *j*′ = *j* - 1, *k*′ = *k* + 1.

Rule 9. *T*′ = *T* \ 〈*j+k*, *I*〉 ∪ 〈*j+k, R*〉, *j*′ = *j, k*′ = *k* + 1. □

### Lemma 4.

*If S*_*j*_*s are all distinct then for any two distinct L*, 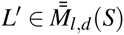, *we have* Γ(*L*) ≠ Γ(*L*^*′*^).

*Proof.* We prove by contradiction. Let *L* = 〈*M, S*_*j,k*_, *T*〉 and *L*^*′*^ = 〈*M, S*_*j*′,*k*′_, *T*′〉 be two distinct elements of 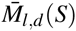. We start with the four strings *O* = *S*_*j,k*_, *O*^*′*^ = *S*_*j*′,*k*′_, *N* = *N*^*′*^ = *M* and align them as follows. The characters common to all four are first aligned. For each *S*_*p*_ present only in one of *O, O′*, we insert *S*_*p*_ in the other and insert *-* in one of *N, N*′, as appropriate. For each 〈*p, t*〉 ∈ *T*, if *t* = *I* we insert a *-* at the corresponding position in *O, O′, N′*. If *t* = *D* we insert a *-* at the corresponding position in *N*. If *t* = *R* we align the corresponding * in *N* with *S*_*p*_ in *O*. We repeat the analogous for 〈*p, t*〉 ∈ *T*′ but making sure that only a single *-* is inserted if both *T, T ′* have an insertion at the same position, or both *T, T ′* have a deletion at the same position.

Without loss of generality, assume *j≤ j′*. If *j < j′* then all of *S*_*j*_, *S*_*j*+1_, …, *S*_*j*′-1_ are either deleted or substituted in *N*. If *S _j,k_* is the not the rightmost *k*-mer of *S* then by Rule 1, *S*_*j*_ cannot be deleted in *N* and hence must be substituted. Then by Rule 3 all of *S*_*j*+1_, …, *S*_*j*′-1_ are also substituted in *N*. Since the leftmost non *-* character in *N*^*′*^ must be *, *S*_*j′,k*′_ must be substituted in *N*^*′*^ because by Rule 7 no insertion is possible just before *S*_*j*′_ in *N*^*′*^. Since *S*_*j*′_ cannot be deleted in *N* by Rule 3, *S*_*j*′_ must be substituted in *N*. This implies there must be another * just after the alignment of *S*_*j*′_ in *N*^*′*^. Since by Rule 8, this * cannot be due to an insertion, *S*_*j*′+1_ must be substituted in *N*^*′*^ which enforces *S*_*j*′+1_ to be substituted in *N*. By repeating this argument, all characters in *N* would be which is not possible. Thus either *j* = *j*^*′*^ or *S _j,k_* is the rightmost *k*-mer of *S*. If *j* ≠ *j*^*′*^ then *S _j,k_* must be rightmost and if *S*_*j*_ is substituted in *N* then a similar argument will show a contradiction. Thus in such a case *S*_*j*_ is deleted in *N* and by Rule 2, each of *S*_*j*+1_, …, *S*_*j*′-1_ is deleted in *N*. By the construction of the alignment there is a at the corresponding positions in *N*^*′*^. Thus in both cases, *j* = *j*^*′*^ and *j < j′*, we have the pattern 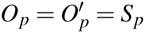 and 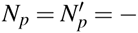 in the alignment for all *p < j′*.

Without loss of generality, we assume *j* + *k ≤ j′* + *k*^*′*^ as we can always interchange *j* + *k* and *j*^*′*^ + *k*^*′*^ in the following argument. If *j* + *k < j′* + *k*^*′*^, let *p* be the leftmost in [*j* + *k* + 1, *′* + *k′ -* 1] such that *S*_*p*_ is deleted in *N*^*′*^. Then by Rule 2, all of *S*_*p*+1_, *S*_*p*+2_, …, S _j′+*k′*-1_ must be deleted in *N*^*′*^. If *p > j* + *k* or no such *p* exists then there is at least one * on the right of the alignment of *S*_*j*+*k-*1_ in *N^I^*. By Rule 9, *S*_*j*+*k-*1_ must be substituted in *N* and hence by Rule 2, *S*_*j*+*k-*1_ must be substituted in *N^I^*, and so on. This will imply all characters is *N* are * which is not possible. Thus *p ≤ j* + *k* and in both cases *j* + *k* = *j^I^* + *k*^*′*^ and *j* + *k < j′* + *k*^*′*^, we have the pattern 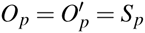 and 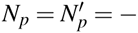 in the alignment for all *p ≥ j* + *k*.

Now let *p* be the leftmost position in the alignment where at least one of the two pairs *O*_*p*_, 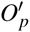 and *N*_*p*_, 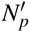 differs. In general, *O*_*p*_, 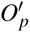 are either both equal to *σ* ∈ Σ or both equal to and *N*_*p*_, 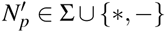. Table 2 shows the 8 possible restricted values for *O*_*p*_, *N*_*p*_, 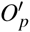, 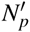 under the assumption of *p*. Table 2 further shows why the cases 1, 2, 3, 5, 6, 8 are not possible. In case 4, to match *N*_*p*_ = * there must be some *p*′ > *p* such that 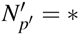. By Rules 2 and 5, *p*′ ≠ *p* + 1. Thus *p*′ > *p* + 1 and 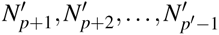 all must be *–*. By Rule 2, these *–*s can not be due to deletions in *S*_*j*′,*k*′_ and hence must be due to insertions in *S*_*j,k*_. Since insertions to both *S*_*j,k*_, *S*_*j*′,*k*′_ at the same position are aligned together, 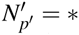 can not be due to an insertion in *S*_*j*′,*k*′_, it must be due a substitution in *S*_*j*′,*k*′_. The corresponding character in *S*_*j,k*_ must be either deleted or substituted. Both are not possible due to Rules 4 and 6 respectively.

**Table 2.** Possible alignments at position *p* where at least one of the pairs *O*_*p*_, 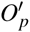 and *N*_*p*_, 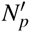 differs

| Case | $O_p$ | $N_p$ | $O'_p$ | $N'_p$ | Comments |
| --- | --- | --- | --- | --- | --- |
| 1 | $\sigma$ | $\sigma$ | $\sigma$ | * | Not possible as there is no other character to match $\sigma$ in $N'$ |
| 2 | $\sigma$ | $\sigma$ | $\sigma$ | — | Same as case 1 |
| 3 | $\sigma$ | * | $\sigma$ | $\sigma$ | Symmetric to case 1 |
| 4 | $\sigma$ | * | $\sigma$ | — | Discussed in details in the text |
| 5 | $\sigma$ | — | $\sigma$ | $\sigma$ | Symmetric to case 2 |
| 6 | $\sigma$ | — | $\sigma$ | * | Symmetric to case 4 |
| 7 | — | * | — | — | Discussed in details in the text |
| 8 | — | — | — | * | Symmetric to case 7 |

In case 7 too, to match *N*_*p*_ = * there must be some *p*′ > *p* such that 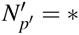. If *p*^*′*^ = *p* + 1 then 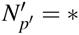 must be due tosome substitution in *S*_*j*′,*k*′_ and the corresponding character in *S*_*j,k*_ must be either deleted or substituted which are not possible by Rules 4 and 6 respectively. Hence *p*′ > *p* + 1 and 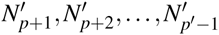 all must be *–*. These *–*s can not be due to deletions in *S*_*j*′,*k*′_ because in that case the corresponding characters in *S*_*j,k*_ must be either deleted or substituted which are not possible due to Rules 4 and 6 respectively. Thus 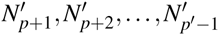 must be due to insertions in *S*_*j,k*_. A similar argument as used in case 4 shows a contradiction in this case too. □

**Algorithm 2:**
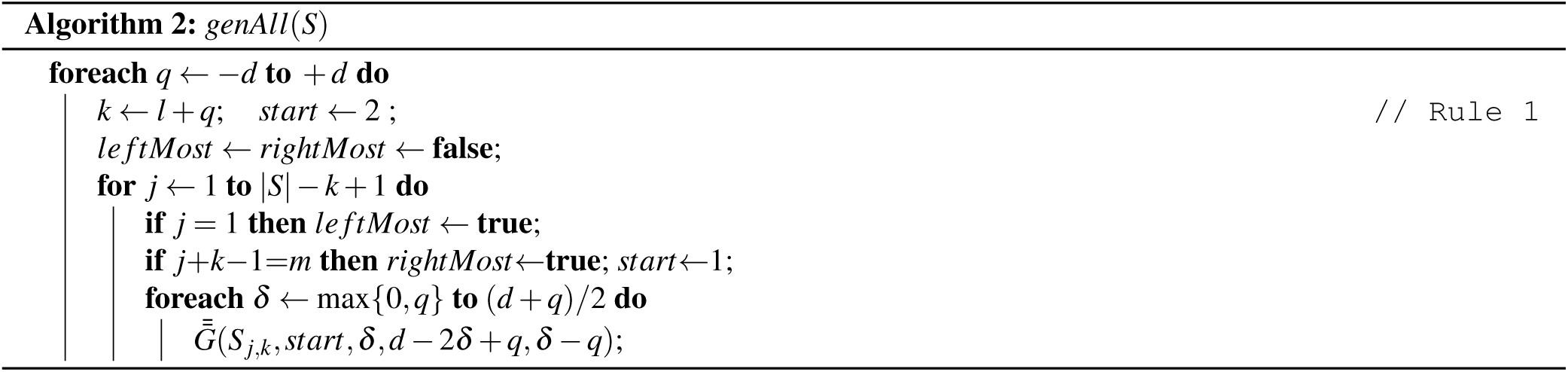
*genAll*(*S*)

**Algorithm 3:**
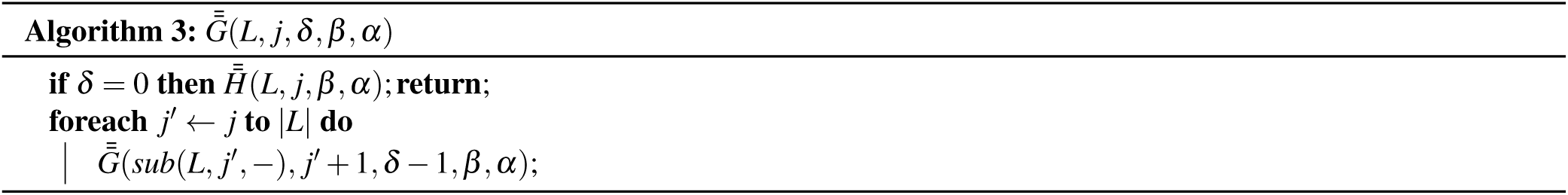
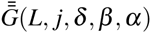

**Algorithm 4:**
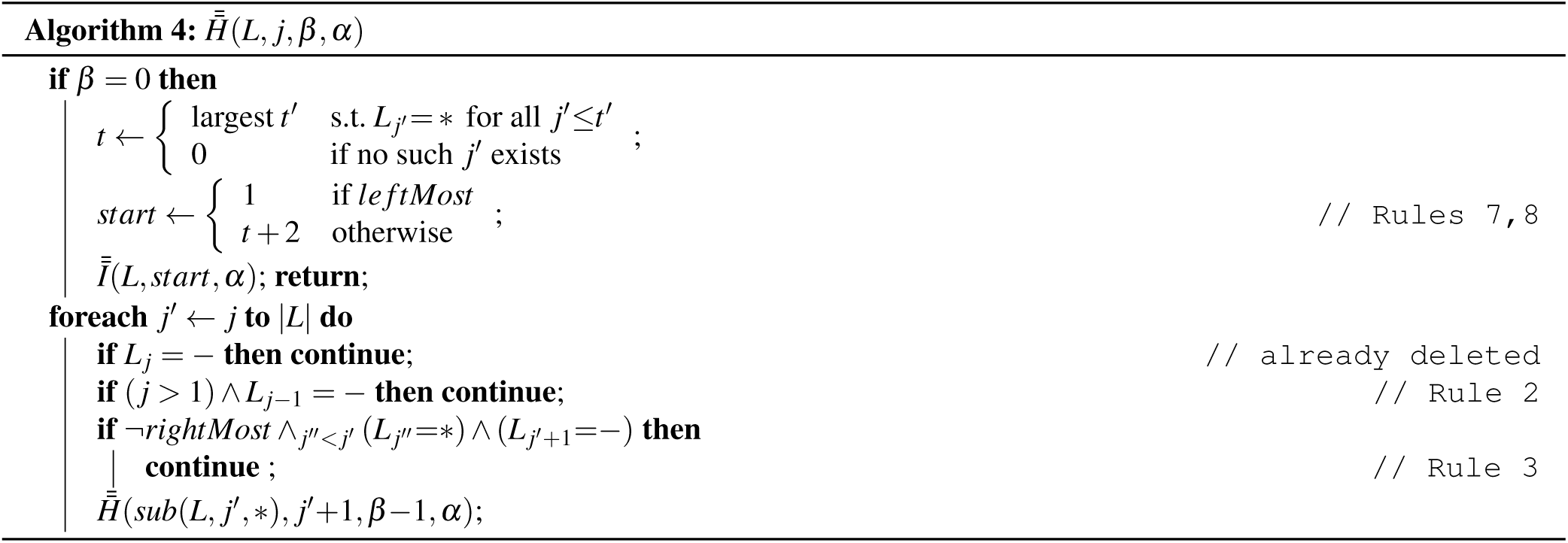
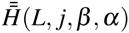

**Algorithm 5:**
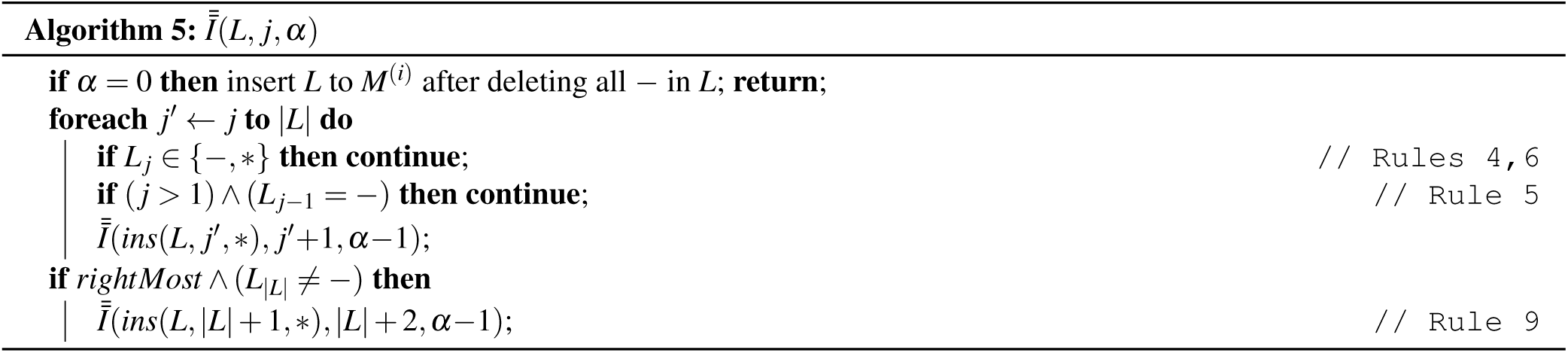
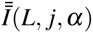

In general the *S*_*j*_s are not distinct. However, as the input strings are random, the repetitions due to repeated characters are limited. Our experimentation shows that on an average each compact motif in 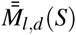 is repeated *c* times where *c* is a real number in [1, 2]. Thus we can safely say that Rules 1-9 make 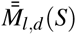 almost duplicate free.

**Implementation:** To track the deleted characters, instead of actually deleting we substitute them by a new symbol — not in Σ*′*. We populate the motif trie *M*^(*i*)^ by calling *genAll*(*S*^(*i*)^) given in Algorithm 2. Rules 1-8 are incorporated in 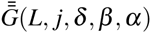, 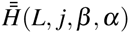 and 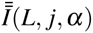 which are shown in Algorithms 3, 4, and 5, respectively.

## Results

We have implemented our sequential algorithm in C++ and evaluated on a Dell Optiplex 7020 desktop with Intel i5-4590 CPU at 3.30GHz and 8GB RAM running Linux Mint 17 (Ubuntu 14.04) OS. We generated random (*l, d*) instances according to Pevzner and Sze^2^ and as described in the introduction. For every (*l, d*) combination we report the average runtime over 5 random instances. We compare our algorithm EMS2 with a modified implementation of the algorithm EMS1^11^ which considered the neighbourhood of only *l*-mers whereas the modified version considers the neighbourhood of all *k*-mers where *l* - *d* ≤ *k* ≤ *l* + *d*.

A comparison between the runtime and the memory usage of the two algorithms are given in Table 3. Our efficient procedure to generate neighbourhood enables our algorithm to solve the instance (13, 4) in less than two hours which EMS1 could not solve even in 3 days. For the instance (13, 4), the memory used by EMS1 is computed using extrapolation (91*(91*/*3)) as EMS1 did not complete in the stipulated time. Note that the factor by which EMS2 takes more memory compared to EMS1 gradually decreases as the instances become harder. Our current implementation of EMS2 stores 4 child pointers in each node of the motif trie corresponding to each edge label *A,C, G, T*. We are working on an implementation in which we need two pointers: one for the leftmost child and one for the immediate right sibling. This is expected to reduce memory usage by about a half for DNA sequences and significantly for protein sequences.

**Table 3.** Comparison between EMS1 and EMS2 on challenging instances. Time is in seconds (s), minutes (m) or hours (h). An empty cell implies the algorithm did not complete in the stipulated 72 hours. †estimated value.

| Instance | Run Time |  | Memory Usage |  |
| --- | --- | --- | --- | --- |
|  | EMS1 | EMS2 | EMS1 | EMS2 |
| (9,2) | 11.9s | 2.6s | 3 MB | 27 MB |
| (11,3) | 35.8m | 1.8m | 91 MB | 474 MB |
| (13,4) | — | 1.2h | 2,760† MB | 7,264 MB |

**Table 4.**
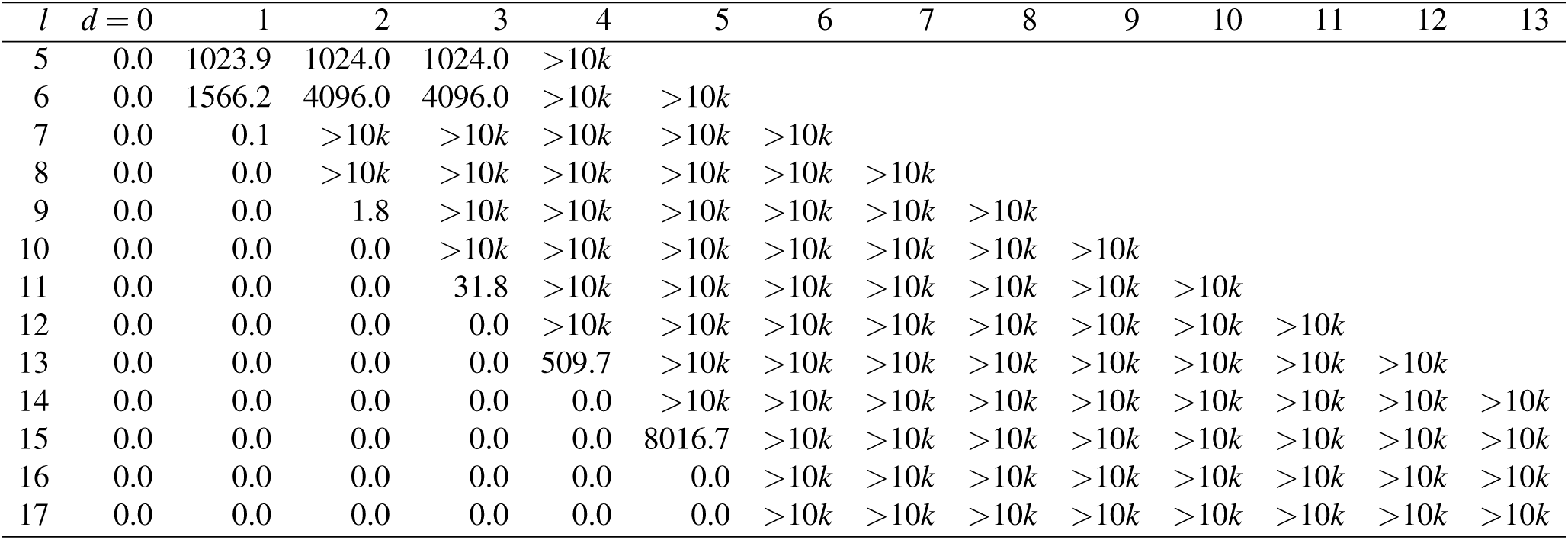
Expected Number of Spurious Motifs in Random Instances for *n* = 20, *m* = 600

## Conclusions

We have presented an efficient algorithm for the EMS problem. Our algorithm is able to efficiently generate neighbourhood using some novel and elegant rules to avoid multiple instances of the same motif being generated in the neighbourhood. We have also proved that these rules are close to the ideal in the sense that the neighbourhood generated by our algorithm is distinct if the characters in the string are distinct. Though this condition is not practical and ideas by Knuth^13^ can be used to generate distinct neighbourhood even when the characters in the string are repeated, nevertheless the rules help because the instances are randomly generated and hence the number of times a *k*-mer appears in any input string is very small. The second reason for the efficiency of our algorithm is the use of a trie based data structure to efficiently and compactly store the motifs.

## Acknowledgments

This work has been supported in part by the NIH grant R01-LM010101.

## Author contributions statement

S.P. and S.R. designed the algorithms, S.P. conducted the experiments, S.P. and S.R. analysed the results, wrote and reviewed the manuscript.

## Additional information

The authors declare no competing financial interest.

## Expected Number of Spurious Motifs

Let *L* be a substring of length *l - q*, 0 ≤ *q* ≤ *d*. If *L* is an occurrence of a fixed motif *M* with *δ* deletions, *α* insertions and *β* substitutions then *α* = *q* + *δ,* 0 *≤ δ ≤* (*d - q*)*/*2, and the number of such *d*-neighbours of *L* is

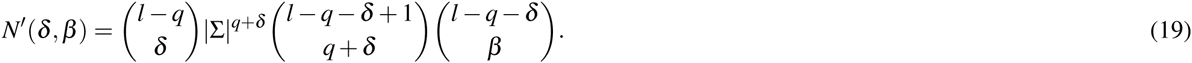

and the probability that *M* is such a neighbour of *L* is

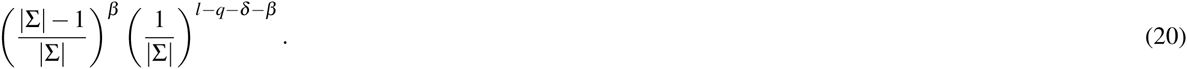

Thus the probability that *L* is an occurrence of *M* is

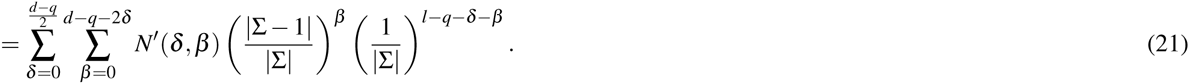

Similarly, if *L* is a substring of length *l* + *q*, 0 *< q ≤ d* the probability that *L* is an occurrence of *M* is

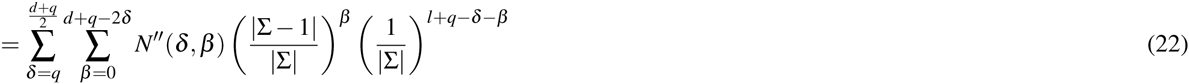

where

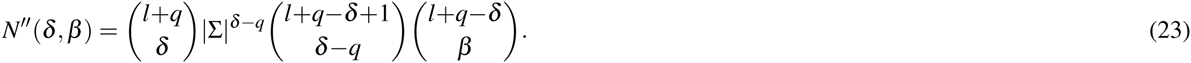

Combining the two cases, for *L* of length *l* + *q*, *-d ≤ q ≤ d*, the probability that *L* is an occurrence of *M* is *P* =

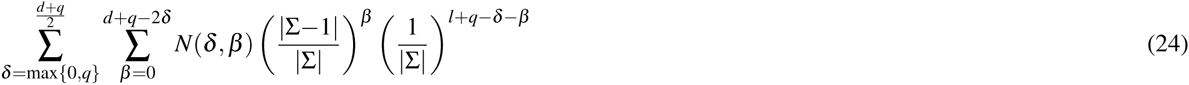

where

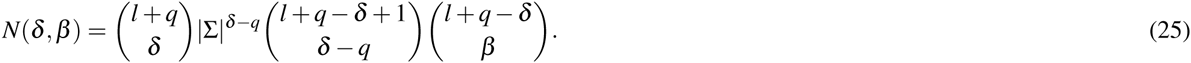

There could be (*m - l - q* + 1) number of (*l* + *q*)-mers of a string of length *m*. The probability that *M* does not occur in the string is

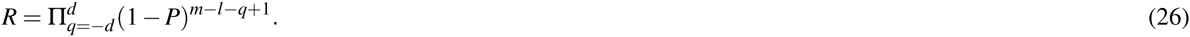

The probability that *M* occurs in each of the input string is (1 *- R*)^*n*^. Since *M* can be any arbitrary motif, the expected number of motifs of length *l* at an edit distance *d* is

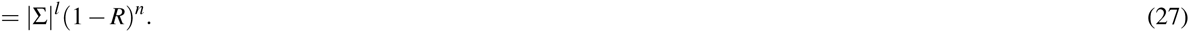

